# Symbiotic nutrient exchange enhances the long-term survival of cassiosomes, the autonomous stinging-cell structures of *Cassiopea*

**DOI:** 10.1101/2023.06.14.544940

**Authors:** Gaëlle Toullec, Niclas Heidelberg Lyndby, Guilhem Banc-Prandi, Claudia Pogoreutz, Cristina Martin Olmos, Anders Meibom, Nils Rädecker

## Abstract

Medusae of the widely distributed and locally invasive upside-down jellyfish *Cassiopea* release autonomous, mobile stinging structures. These so-called cassiosomes are a major contributor to ‘contactless’ stinging incidents in (sub-)tropical shallow waters. While the presence of endosymbiotic dinoflagellates in cassiosomes has previously been observed, their potential contribution to the metabolism and long-term survival of cassiosomes is unknown. Combining stable isotope labeling and correlative SEM and NanoSIMS imaging with a long-term *in vitro* experiment, this study reveals a mutualistic symbiosis based on nutritional exchanges in dinoflagellate-bearing cassiosomes. We were able to show that organic carbon input from the dinoflagellates fuels the metabolism of the host tissue and enables anabolic nitrogen assimilation. Thanks to this symbiotic nutrient exchange, cassiosomes showed enhanced survival in the light compared to dark conditions for at least one month *in vitro*. Overall, this study demonstrates that cassiosomes, in analogy with *Cassiopea* medusae, are photosymbiotic holobionts. Cassiosomes thus promise to be a powerful new miniaturized model system for in-depth ultrastructural and molecular investigation of cnidarian photosymbioses.

## Introduction

Jellyfish (scyphomedusae) blooms can have significant impacts on marine ecosystems and the human communities depending on them. Beyond their role in the marine food web, in biogeochemical cycling, and in fishery, jellyfish blooms also lead to increases in sting-related injuries among swimmers. Because of this sting threat, jellyfish blooms thus have a strong negative effect on coastal tourism (1–4). Jellyfish blooms have been linked to the rise of sea surface temperatures and other human disturbances, such as eutrophication and overfishing (1, 5). The frequency and extent of these blooms have thus been predicted to locally increase or oscillate in the future (5–8).

The upside-down jellyfish Cassiopea (Scyphozoa, Rhizostomae) have recently been reported as newly introduced and locally invasive in numerous localities (9–11). Due to their relatively high heat tolerance and trophic plasticity, their population density and geographic expansion are only expected to increase further (12–15). Like all cnidarians, *Cassiopea* medusae harbor specialized stinging cells called nematocytes that play an important role in predator defense and prey capture. While *Cassiopea* stings are often considered mild, Muffet and al. (16) recently highlighted their potential severity and a lack of public awareness regarding their threat. Jellyfish stings by direct contact are well known, but ‘contactless’ stings without direct physical contact with the animal have also been reported (16). Among contactless stinging mechanisms, the release of cassiosomes (i.e., autonomous, stinging and often motile tissue structures) has been recently described in several rhizostome medusae, including some *Cassiopea* species (17, 18). Interestingly, the cassiosomes from *C. xamachana, C. ornata* and two mastigiidae medusae host phototrophic dinoflagellates of the Symbiodiniaceae family, a group known to form intimate symbiotic relationships with a diversity of cnidarians, such as corals and sea anemones (17–21). While *Cassiopea* polyps and medusae benefit strongly from organic carbon input from their dinoflagellate symbionts and are now well-established model systems for the cnidarian-*Symbiodiniaceae* symbiosis (22–26), the contribution of dinoflagellates to the metabolism and survival of cassiosomes remains unknown. Disentangling the metabolic activity and survival capacity of cassiosomes is therefore a key step to understand and predict the stinging threat represented by these cassiosomes in the marine environment. Furthermore, if indeed cassiosome symbiotic nutrient cycling is analogous to that of their medusae of origin, these small, easily and year-long collectable tissue structures have the potential to become an important miniaturized model system for a variety of studies of photosymbiotic interactions.

In this study, we first describe the ultrastructure of *C. andromeda* (Forskål, 1775) cassiosomes with a workflow including high-pressure freezing and CryoSEM imaging. We then test the metabolic activity and potential nutritional exchange between cassiosome tissues and their dinoflagellates by stable isotopic labeling and correlative SEM and NanoSIMS imaging. Finally, we assessed the contribution of dinoflagellates photosynthates to cassiosomes survival by maintaining cohorts of cassiosomes on either a 12h:12h day-night light cycle or in complete darkness in a two-month-long *in vitro* experiment (**Figure 1**).

**Figure 1:**
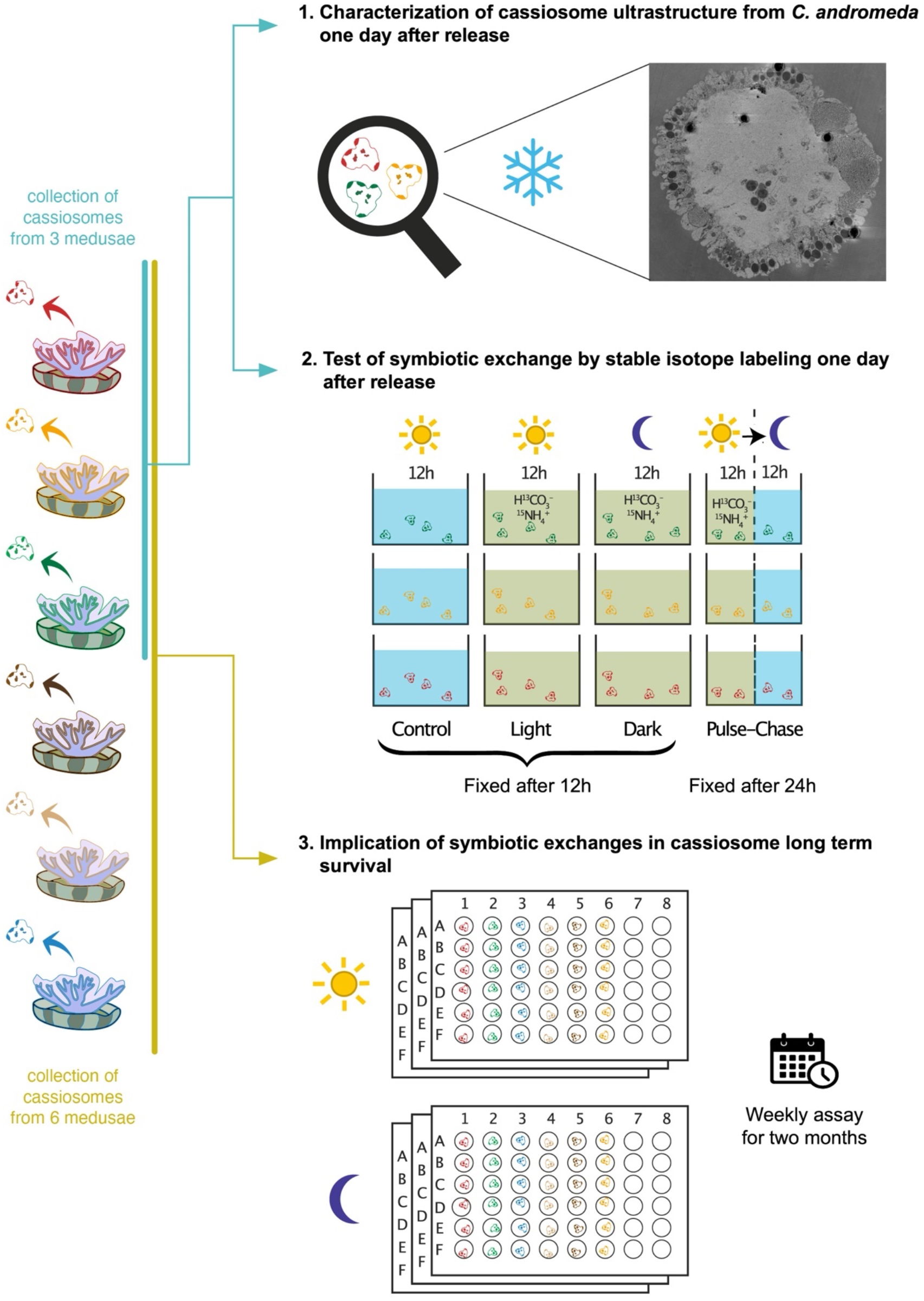
Schematic illustration of the study design composed of three experiments. The colors of cassiosomes in the individual experiments indicate that they were originally collected from different medusae. The number of cassiosomes per beaker in the second experiment is for illustrative purposes only and not a quantitative representation of the actual experiment (cf. Materials and Methods).

## Materials and methods

### Animal husbandry and cassiosome collection

Adult *Cassiopea* medusae were acquired from De Jong Marinelife in the Netherlands. Amplification and sequencing of fragments of the COI (mitochondrial cytochrome oxidase subunit) regions from three individuals identified the species as *Cassiopea andromeda* (data not shown). In the 200 L culture aquarium, the medusae were maintained in artificial seawater (ASW) prepared from sea salts (Reef Salt, Aquaforest) at a constant salinity of 35 ppt and temperature of 25°C, illuminated with approximately 100 μmol photons m^−2^ s^−1^ (400-700 nm) from LED lights on a 12h:12h day:night cycle. The medusae were fed *ad libitum* two to three times a week with freshly hatched *Artemia salina* nauplii.

For the experiments, cassiosomes were collected from individual medusae of approximately 5 cm in diameter. Animals were gently sprayed with a jet of ASW in a small beaker to cause the release of cassiosomes. 100 mL of ASW containing cassiosomes was collected from each animal and placed overnight in an incubator at 25 °C on a 12h:12h day:night cycle, in order to separate the sinking cassiosomes from the floating mucus prior to their use in any of the experiments (17).

### Characterization of cassiosomes ultrastructure by cryo-SEM

In order to characterize the ultrastructure of the cassiosomes in their most pristine condition, cassiosomes were fixed, prepared and imaged using a fully cryogenic workflow (27–29).

The day following their release, cassiosomes were collected by gentle pipetting from the bottom of a petri dish (thus avoiding the floating mucus) using a stereomicroscope, transferred into a 1.5 ml tube and concentrated by centrifugation at 425 x g for 2 min. The method used for pristine cryopreservation of the cassiosomes was high-pressure freezing (HPF). HPF delivers synchronized pressurization and cooling of small samples (< 200μm thick) with liquid nitrogen within 20 ms, thereby avoiding any nucleation of ice crystals that would damage the tissue ultrastructure (27). For this, the pellet of cassiosomes was resuspended in a small volume of the cryoprotectant 20 % dextran 40 (prepared in 35 ppt ASW, Sigma D-1662, US). A small amount of the resuspended cassiosomes were pipetted into an Aucoated Cu-carrier and high-pressure frozen using a Leica EM ICE high-pressure freezer (Leica Microsystems, Germany). Cryopreserved samples were cryo-planed with a diamond trim knife (DiATOME, Switzerland) using a UC7 ultramicrotome (Leica Microsystems, Germany) at -110 °C, and transferred to a Leica EM ACE 600 (Leica Microsystems, Germany) for a two-step process. First, freeze etching was performed in order to eliminate any surface ice contamination deposited after trimming and to create morphological contrast between cellular components (28). For this, the sample was warmed up from -150 °C to -93 °C with a ramp of 3 °C min^-1^, then held at -93 °C for 2 min and brought back to -150 °C at a rate of 3 °C min ^-1^. Second, a 3 nm platinum layer was deposited by e-beam evaporation at -150 °C to minimize surface charging during subsequent cryo-SEM imaging. Finally, the samples were transferred and imaged by cryo-scanning electron microscopy (SEM, GeminiSEM 500, Zeiss, Germany; 1.7 kV, aperture size of 10 μm, and a working distance of 3.4 mm) with an Inlens detector (Zeiss, Germany). Cryo-SEM images were adjusted in contrast and brightness, as well as artificially colored for optimized visualization of the structures using Photoshop software (Adobe Photoshop 2023, version 24.3.0).

### Stable isotope labeling experiment

In order to investigate the uptake and exchange of nutrients between the cassiosomes and their Symbiodiniaceae symbionts, a stable isotope labeling experiment was performed using cassiosomes one day after their release from three adult medusae (i.e., three independent biological replicates in total).

The day before the labeling experiment, filtered ASW was depleted of any dissolved organic carbon by acidification with HCl (4 M) to a pH < 3, and maintained under constant air bubbling for at least 4 h. This ASW was then labeled with ^13^C-bicarbonate (#372382, Sigma-Aldrich, USA) to a final concentration of 3 mM. Finally, the pH of the solution was raised again to 8.1 with 1 M NaOH solution and labeled with 15N-ammonium-chloride (#299251, Sigma-Aldrich, USA) to a final concentration of 3 μM. After thorough homogenization, the labeled ASW and freshly prepared unlabelled ASW were prewarmed and maintained at 25 °C overnight.

On the morning of the experiment, floating mucus was removed by pipetting off 20 mL of water from the surface of the three beakers containing the cassiosomes. The remaining content in each of the beakers was gently mixed, split into four equal fractions of 20 mL and concentrated by filtration through a 40 μm cell strainer (Corning, USA). The four fractions of each cassiosome sample were resuspended in 40 mL of labeled or unlabeled ASW accordingly.

Subsequently, cassiosomes collected from each medusa (n = 3) were subjected to four different experimental conditions: light, dark, pulse-chase, and control (**Figure 1**). In order to assess the nutrient assimilation by the cassiosomes and their endosymbiont algae with or without photosynthesis, incubations of 12 h in labeled ASW were performed in light and darkness, respectively. In addition, in order to assess potential relocations over time of the nutrients assimilated during the light period, a pulse-chase experiment was carried out consisting of 12 h incubation in labeled ASW in light followed by 12 h in unlabeled ASW in darkness. Finally, the remaining cassiosome batches were maintained in unlabeled ASW in the light for 12 h to generate unlabeled control samples with natural isotopic composition of both cassiosomes and algae.

The labeling incubation was performed in glass beakers maintained at 25 °C in a 15 L water bath equipped with a circulation pump and a heater. During the incubation, the samples were illuminated for 12 h with LED lights (Viparspectra V165, USA) providing approximately 100 μmol photons m^−2^ s^−1^ (400-700 nm) with the exception of the dark condition, which was maintained in constant darkness with aluminum foil. To ensure constant isotopic concentrations in the labeled incubation water, the water of each beaker was gently mixed every 2 hours, and half of the volume of labeled or unlabeled ASW was replaced in each sample every 4 hours. At the end of the 12 h incubation, all samples were gently concentrated by filtration using a cell strainer (40 μm mesh size), and the samples corresponding to light, dark, and control conditions were resuspended in 3 mL of fixative solution (4 % paraformaldehyde, 2.5 % glutaraldehyde in 0.1 M Sorensen’s buffer, 9 % sucrose) for 16 h before further processing. The samples subjected to a pulse-chase were resuspended in unlabeled seawater, and incubated for 12 more hours in the dark. At the end of this chase period, the cassiosomes were filtered and chemically fixed as previously described for 4 hours.

### Sample preparation for correlative SEM-NanoSIMS imaging and analysis

After fixation, all the samples from the isotope labeling experiment were prepared for correlative SEM and NanoSIMS imaging.

Each of the 12 cassiosome samples (four treatments and three source medusae) was split into two aliquots in 1.5 mL Eppendorf tubes and rinsed twice to remove the fixative (centrifugation at 425 x g for 5 min and rinsed by resuspension in 0.1 M Sorensen’s buffer). To preserve the lipid fraction of the samples, a post-fixation was performed for 1 h with osmium tetroxide (OsO4 1 %, 1.5 % potassium hexacyanoferrate II in 0.1 M Sorensen phosphate buffer) under constant agitation, and rinsed by centrifugation and resuspension in milli-Q water under constant agitation for 15 min. After another centrifugation cycle, the samples were pre-embedded in agarose to avoid the loss of cassiosomes in the subsequent steps. 20 μL of the cassiosome pellets were transferred into 400 μL polyethylene microtubes (#391178, Milian) prefilled with 200 μL of 2 % liquid agarose at 40 °C, and immediately centrifuged at 20800 x *g* for less than 1 min. After curing on ice for 5 min, the tubes were cut open and the agarose-embedded pellets of cassiosomes were dissected into pieces of approximately 1 mm^3^. Using a tissue processor (Leica Microsystems, Germany), the samples were then subjected to a serial dehydration in ethanol (30 %, 70 % and 100 % ethanol in Milli-Q water), to facilitate a progressive Spurr resin infiltration of the samples (30 %, 70 % and 100 % Spurr resin in absolute ethanol). Once infiltrated, the samples were transferred into molds filled with 100 % Spurr resin and cured at 60 °C for 48 hours. Semi-thin sections (200 nm) of the samples were cut from the resin blocks using an Ultracut S microtome (Leica Microsystems, Germany) and a diamond knife. These sections were then transferred to clean glow-discharged glass slides (for NanoSIMS analysis) or silicon wafers (for correlative SEM and NanoSIMS imaging). In order to add contrast and to visualize the subcellular structures of the cassiosomes and the algae, the sections on silicon wafers were post-stained with 1 % uranyl acetate and Reynolds Lead Citrate before imaging by scanning electron microscopy (SEM, Gemini 500, Zeiss, Germany; 3 kV, aperture size of 30 μm, and a working distance of 2.9 to 2.3 mm) with an energy selective backscatter detector (EsB, grid of 130 V; Zeiss, Germany). Prior to NanoSIMS imaging (30), sections were sputter coated with a 12 nm gold layer (using a Leica EM SCD050 gold coater). In the NanoSIMS, the pre-sputtered samples were bombarded with a Cs^+^ primary ion beam at 16 keV with a current of around 2 pA, focused to a spot size of ca. 150 nm. For each image, this beam was rastered over an area of 40 x 40 μm with a resolution of 256 × 256 pixels and a dwelling time of 5000 μs per pixel for five consecutive layers. The secondary ions ^12^C^12^C^-, 12^C^13^C^-, 12^C^14^N^-, 12^C^15^N^-^ were counted individually in electron multiplier detectors at a mass resolution power of around 9000 (Cameca definition), which resolves potential interferences in the mass-spectrum. The resulting isotopic maps were analyzed using L’Image (v.10-15-2021, developed by Dr. Larry Nittler, Arizona State University). Images were drift corrected and regions of interest (ROIs) were drawn around the different compartments, i.e., dinoflagellates, amoebocytes (excluding the dinoflagellates), and the cassiosome epidermis. For each ROI, the isotopic ratio enrichments established through the ratios ^12^C^13^C^-^/^12^C2-and 15N^12^C^−^/^14^N^12^C^−^ were quantified against a control sample with natural isotopic compositions prepared and analyzed in an identical manner. Isotope enrichments are expressed in the delta-notation as followed:

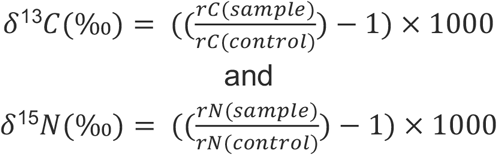

where rC(sample) and rC(control) are the count ratios of ^12^C^13^C^-^/^12^C2-in the sample and in the unlabeled control, respectively. rN(sample) and rN(control) are the count ratios of 15N^12^C^−^/^14^N^12^C^−^ in the sample and the control, respectively. A compartment was only considered to be isotopically enriched if its average delta-value was more than three standard deviations above the average ratio measured in similar compartments in the unlabeled sample.

### Long-term survival experiment

In order to investigate the contribution of symbiont photosynthesis to the long-term survival of these autonomous structures, cassiosomes from 6 adult medusae were maintained in light or dark conditions for two months (**Figure 1**).

One day after being released, cassiosomes (36 per medusa) were distributed (one per well) over six sterile flat-bottom 48 well plates (Costar 3548, Corning, USA) in 1.5 mL of filter-sterilized ASW (filtered through 0.22 μm pore size) at a salinity of approximately 35 ppt. Cassiosomes selected for the experiment were individually observed under a stereomicroscope equipped with a blue light and GFP filter (M165 C, Leica Microsystems, Germany) to verify the motility and presence of pigmented algal symbionts by fluorescence. Each plate was then placed into a humid chamber (sealed in a transparent plastic bag containing wet tissue paper to avoid evaporation). All six humid chambers containing the cassiosomes in 48 well plates were maintained for two months in an incubator at a constant temperature of 25 °C. Three replicate chambers were maintained on a 12 h:12 h day:night light cycle at approximately 100 μmol photons m^−2^ s^−1^ (400-700 nm), and three were maintained in constant darkness (wrapped in aluminum foil and kept in a separate dark compartment in the incubator). The presence/absence of each of the 216 cassiosomes was individually assessed under the binocular on a weekly basis for a duration of 8 weeks, as an estimation of cassiosome survival. A total of 66 % of the ASW in each well was carefully replaced by pipetting to avoid disturbing the cassiosomes.

### Statistical analysis

All statistical analyses were performed in R (version 4.2.0, (31). The difference in isotopic enrichment between experimental conditions was analyzed using a linear mixed model (LMM) with the animal medusa as a random variable. This analysis was followed by a Tukey’s Honestly Significant Differences (HSD) post hoc comparison. The overall impact of light and time on the cassiosome estimated survival was analyzed for the linear phase of the data (from day 0 to 35) using a linear mixed model (LMM) with the medusa of origin as a random variable. The difference between light treatments on each day was then analyzed by a pairwise t-test with a subsequent Bonferroni correction of the p-values.

## Results

### Description of the Cassiopea andromeda cassiosome ultrastructure by light microscopy and cryo-SEM

The cassiosomes collected from *Cassiopea andromeda* showed considerable variation in size and shape, and exhibited motility that can be attributed to the presence and movement of cilia. Most, but not all, of the collected cassiosomes harbored Symbiodiniaceae.

Cryogenic imaging permitted the observation of ultrastructural features of the cassiosomes in their most pristine condition (**Figure 2**). Overall, the cassiosomes consisted of an external epidermal cell layer surrounding a ‘core’ of mesoglea. The epidermal cell layer contained a high density of nematocytes, often grouped in clusters (**Figure 2C,E)**. Inside the mesoglea core, amoebocyte cells were present, many of which were observed to host dinoflagellates (**Figure 2C-D**).

**Figure 2:**
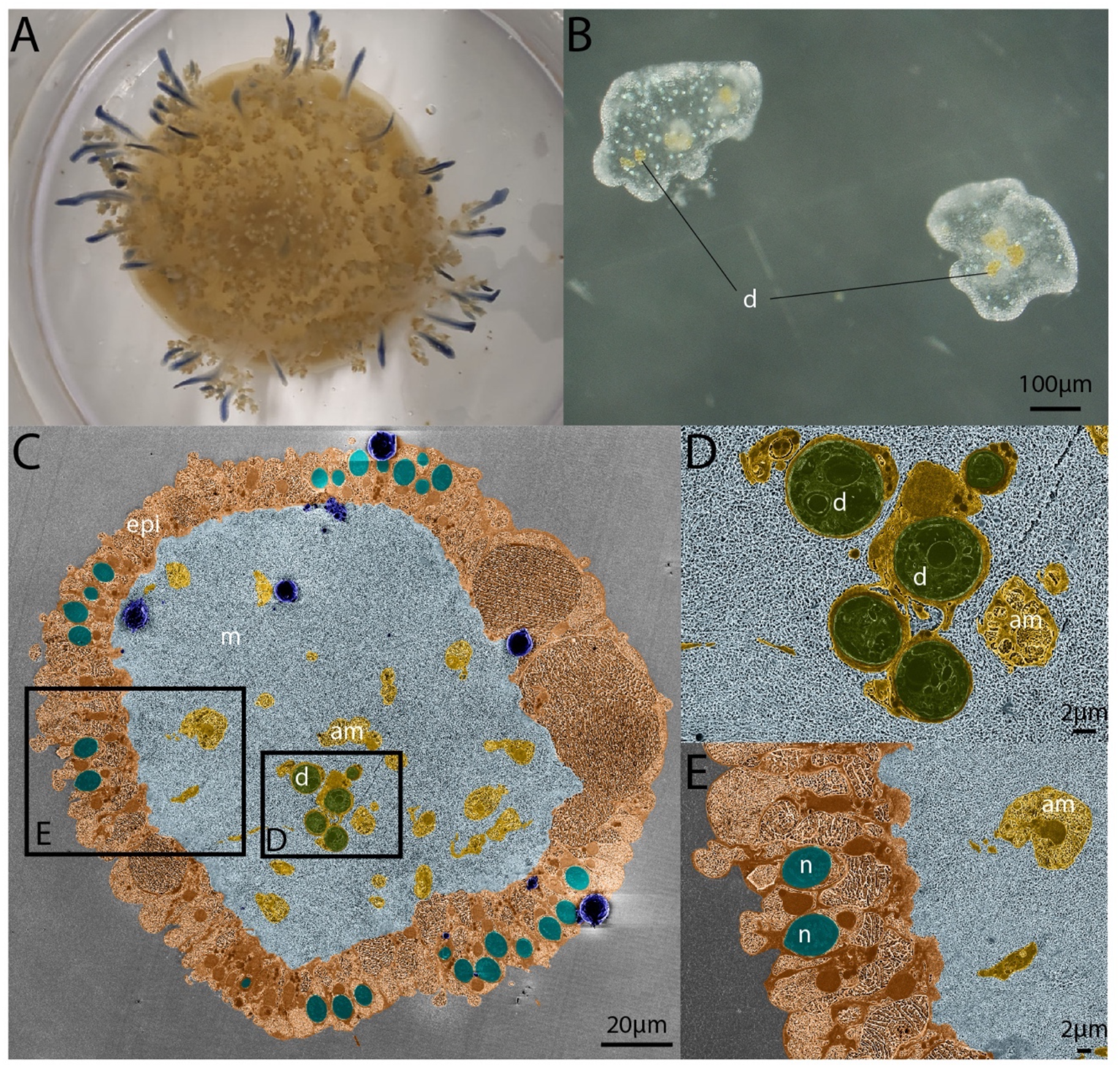
Appearance and ultrastructure of cryopreserved cassiosomes. **A** Adult Cassiopea andromeda medusae (approximately 5 cm bell diameter). **B** Appearance of freshly collected cassiosomes in the stereomicroscope. **C** Cross-section of a representative cassiosome harboring dinoflagellates imaged by cryoSEM. **D** Ultrastructural details of amoebocytes hosting dinoflagellates and **E** of the cassiosome epithelium harboring nematocytes in a representative cassiosome imaged by cryoSEM. CryoSEM images are artificially colored for better visualization of cassiosomes features: d: dinoflagellate (in green), epi: cassiosome epithelium (in orange), m: mesoglea (in light blue), am: amoebocyte (in yellow), n: nematocyst (in turquoise). Ice crystal surface contamination is highlighted in dark blue.

### Nutrient assimilation and exchange in the cassiosomes and their algal symbionts

The correlative SEM-NanoSIMS analysis of the ^13^C-bicarbonate and ^15^N-ammonium labeling experiment revealed active and light-dependent assimilation and translocation of inorganic nutrients within the cassiosome-algal symbiosis (**Figure 3**).

**Figure 3:**
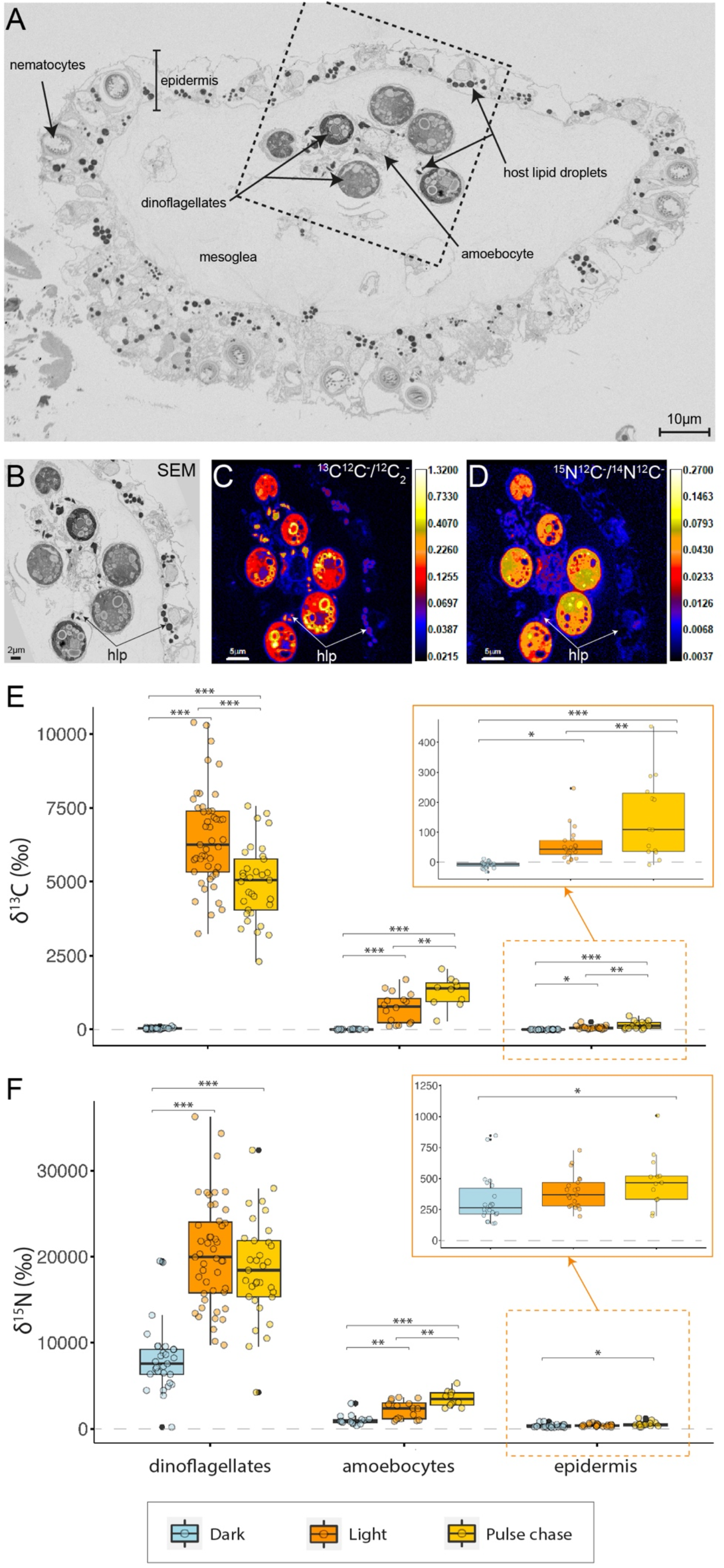
Assimilation of inorganic carbon and nitrogen and translocation of their metabolic derivatives within the cassiosome-algal symbiosis. SEM image of a cross-section of a representative resin-embedded cassiosome (**A**) illustrating its cellular organization. Correlative SEM (**B**) and NanoSIMS (**C, D**) imaging showing the subcellular localization of ^13^C assimilated from ^13^C-bicarbonate (**C**) and ^15^N assimilated from ^15^N-ammonium (**D**) in light (NanoSIMS images are shown with isotope ratios expressed in logarithmic color scale). hlp: host lipid droplets. Quantification of ^13^C enrichment (**E**) and ^15^N enrichment (**F**) in cassiosome compartments (dinoflagellates, amoebocytes and epidermis) under the different experimental conditions. Asterisks indicate significant differences between treatments (* p < 0.050, ** p < 0.010, *** p < 0.001).

After the 12 h incubation in the light, the three measured cassiosome compartments (i.e., dinoflagellates, amoebocytes and epidermis) were significantly enriched in ^13^C (**Figure 3B,C,E**). This ^13^C enrichment was particularly apparent in the pyrenoid and starch granules of the dinoflagellates and in the abundant lipid droplets (dark intracellular bodies stained by osmium in SEM images) present in the amoebocytes and epidermis (**Figure 3B,C**). Similarly, all compartments were enriched in ^15^N (**Figure 3F**). The ^15^N enrichment was distributed quasi-homogeneously within the dinoflagellate cells, the amoebocytes, and the epidermis cells, respectively (**Figure 3B,D**).

The absence of light during the 12 h incubation strongly impacted nutrient assimilation in the cassiosomes (**Figure 3E,F**). In the dark, the ^13^C enrichment was undetectable (below the enrichment threshold) in the dinoflagellates, amoebocytes and epidermis (Tukey’s HSD, *p* <0.001 for dinoflagellates and amoebocytes, *p* = 0.022 for the epidermis, **Figure 3E**). The ^15^N enrichment was also significantly lower in the dark compared to the light condition, specifically by 58 % in the dinoflagellates (Tukey’s HSD, *p* < 0.001), 54 % in the amoebocytes (Tukey’s HSD, *p* = 0.003) and 19 % (albeit not significantly) in the epidermis (Tukey’s HSD, *p* = 0.273, **Figure 3F**).

Finally, the light pulse followed by a dark chase period (pulse-chase condition) revealed the temporal cascade of nutrient assimilation and translocation in the symbiosis. Compared to the light condition (i.e., pulse without a chase period), the ^13^C enrichment in the dinoflagellates decreased significantly by 25 % over the subsequent 12 h dark period (Tukey’s HSD, *p* < 0.001). In contrast, the ^13^C enrichment increased by 67 % in the amoebocytes (Tukey’s HSD, *p* = 0.004) and by 145 % in the epidermis (Tukey’s HSD, *p* = 0.003, **Figure 3E**). In addition, while the ^15^N enrichment remained overall similar following the 12 h dark chase in the dinoflagellates (Tukey’s HSD, *p* = 0.828), it experienced an increase of 56 % in the amoebocytes (Tukey’s HSD, *p* = 0.002) and 36 % epidermis (Tukey’s HSD, *p* = 0.141, **Figure 3F**) after the next 12 h dark period, when compared with the light condition.

### Light enhances cassiosome survival

The two months long culture experiment illustrated the impact of light availability on symbiont-bearing cassiosome survival *in vitro* (**Figure 4**).

**Figure 4:**
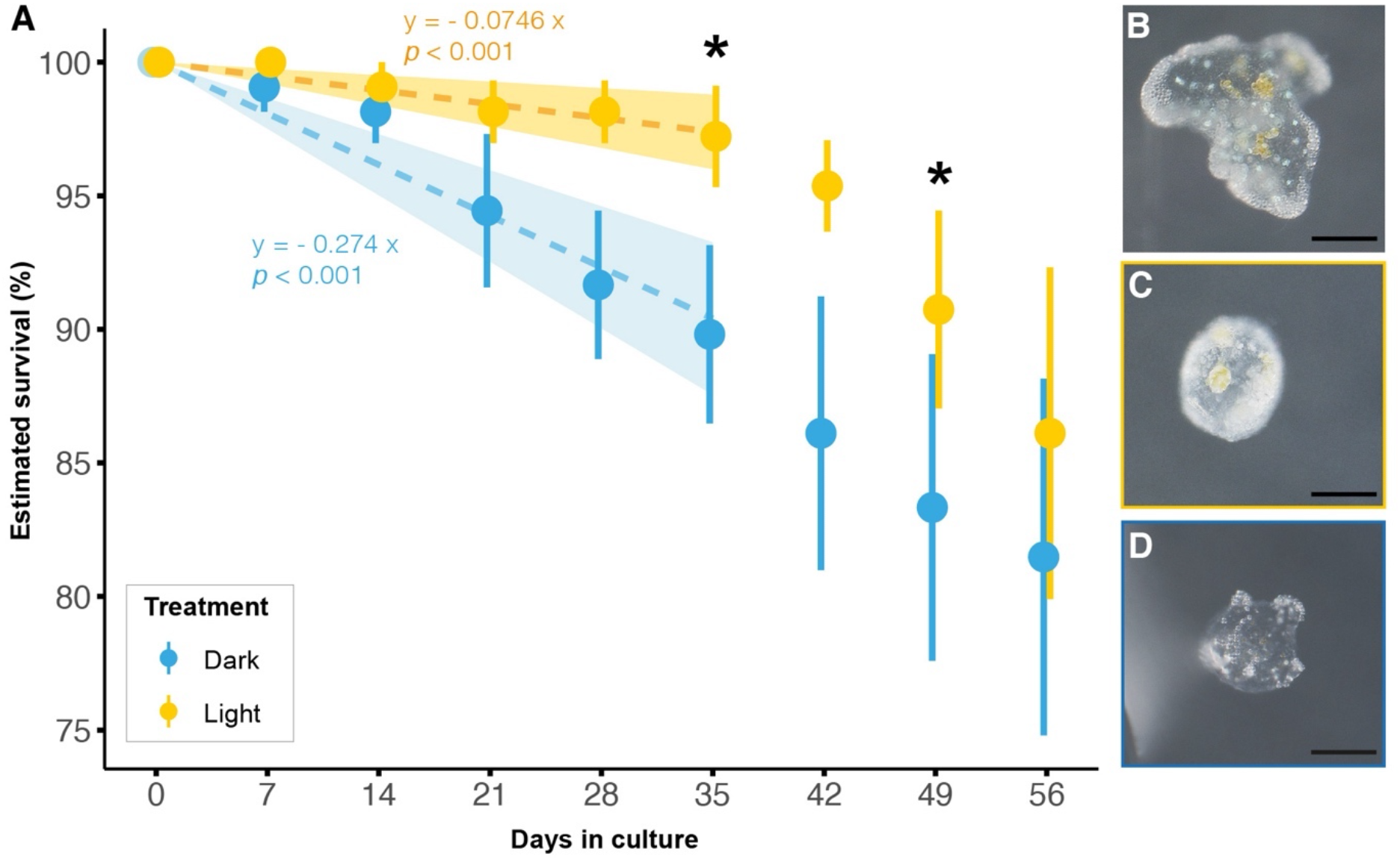
Influence of light treatments on cassiosome survival. **A** Estimated survival of cassiosomes over time in vitro, under a 12 h: 12 h light: dark cycle (yellow) or in constant darkness (blue). Filled circles and error bars indicate the mean ± SE. n = 108 cassiosomes were used per condition, which included cassiosomes from 6 different medusae. Asterisks indicate significant differences between treatments (* p < 0.050) at a given time point. Images of chemically-fixed cassiosomes before (**B**) or after being maintained in light (**C**) or dark (**D**) conditions for 2 months (scale bar = 100 μm).

Overall, the interaction of light treatment and time had a significant effect on cassiosome survival (LMM, *X*^*2*^ = 13.46, *p* < 0.001, **Figure 4A**) during the first 35 first days of the experiment. During this linear phase of the cassiosome decline, the rate of disappearance of cassisomes in the dark was 3.6 fold higher than in the light. Over time, this led to a significant difference in survival between treatments (pairwise t-tests, *p-adjusted* = 0.043, 0.025 for day 35 and 49 respectively, **Figure 4A**). After day 35, the rate of disappearance of cassiosomes in the light increased and exceeded the one in the dark condition.

Of note, more than 80 % of the cassiosomes were still present in both conditions at the end of this two-month experiment, but their appearance changed. The cassiosomes experienced a strong reduction in size, often losing their characteristic ‘pop-corn’-like shape, and became featureless (**Figure 4B,C,D**). While no quantification has been performed, the dinoflagellates in most of the cassiosomes of the light condition were still visible at the end of the experiment while most of the cassiosomes kept in the dark condition showed no chlorophyll fluorescence upon scrutinization in the stereomicroscope.

## Discussion

Global warming and local anthropogenic stressors, such as overfishing and eutrophication, have been linked to recent local increases in jellyfish population density, spatial distribution, and stinging threat (1, 5). Members of the upside-down jellyfish *Cassiopea spp*. have been described as particularly invasive in several tropical and subtropical regions around the world in recent years (9–11). Some *Cassiopea* species and other members of the Rhizostomeae order have been shown to release stinging, autonomous and often motile tissue structures called cassiosomes. These cassiosomes are likely a major contributor to the ‘contactless’ stinging phenomenon (16, 17). Here, we provided the first report, to our knowledge, on the existence and ultrastructure of cassiosomes in *Cassiopea andromeda*. In addition, we showed that dinoflagellates act as beneficial symbionts within the cassiosome by fueling their host’s metabolism with photosynthates and thus prolonging their autonomous life span.

### *The ultrastructure of cassiosomes from* Cassiopea andromeda

The ultrastructure of the cassiosomes of *C. andromeda* resembles those of previously described from *C. xamachana* (17) and *C. ornata* (18). The ability of *C. andromeda*, another rhizostome medusa species, to also produce cassiosomes supports the idea of cassiosomes being an ubiquitous evolutionary feature of the Rhizostomeae order (17).

*C. andromeda*’s cassiosomes were composed of an external epidermis containing a high number of nematocytes, and a core of mesoglea harboring amoebocytes that frequently hosted dinoflagellates (**Figure 2**). This overall cellular organization is similar to the one previously described for cassiosomes from *C. xamachana* (17).

### Algal symbionts contribute photosynthates to cassiosome metabolism

One day after their release, the cassiosomes incubated with ^13^C-bicarbonate and ^15^N-ammonium in light showed strong enrichments in ^13^C and ^15^N in cassiosome cells (amoebocytes and epidermis) and in their dinoflagellates (**Figure 3**). This demonstrates that cassiosomes are anabolically active and are able to assimilate nutrients from the seawater, even after the separation from the medusa.

In the dark, the disappearance of ^13^C enrichment (**Figure 3E**) indicates that the fixation of inorganic carbon from the seawater is primarily driven by dinoflagellate photosynthesis in a light-dependent manner. The associated drop in ^15^N enrichment in the cassiosome cells (**Figure 3F**) is likely caused by a reduction in carbon availability in the cassiosomes in the absence of algal photosynthesis. Indeed, cnidarian ammonium assimilation requires the availability of excess carbon backbones in the TCA cycle for amino acid synthesis.

Consistently with this, following the 12 h pulse labeling in the light, a 12 h dark chase period with unlabeled ASW caused a decrease in ^13^C enrichment in the dinoflagellates and an increase in ^13^C enrichment in amoebocytes and epidermis (**Figure 3E**). This indicates that an active transfer of photosynthetically fixed carbon took place from dinoflagellates to the cassiosome tissues. This increase of ^13^C enrichment was associated with an increase in ^15^N assimilation in the cassiosome tissue (**Figure 3F**), again best ascribed to organic carbon availability in the cassiosome tissue: as carbon availability increases with time in the cassiosome tissue through the translocation process, the cassiosome cells are able to anabolically assimilate more ammonium.

In conclusion, these results demonstrate that cassiosomes maintain an active metabolism that is supported by symbiotic nutrient exchange with their associated dinoflagellates. The algal symbionts fuel and shape the cassiosome metabolism with photosynthetically fixed carbon, in a manner highly similar to the symbiotic interactions observed in the intact medusa (22) and in other marine photosymbioses (32–37).

### *Symbiotic nutrient input supports the long-term survival of cassiosomes* in vitro

The positive impact of light and associated photosynthetic input from the symbiont to cassiosome survival was reflected in a significantly higher estimated survival rate during the first five weeks of the *in vitro* experiment, compared with the dark condition **(Figure 4)**. While light (and associated algal photosynthesis) enhanced cassiosome survival until day 35 compared to the dark condition, this advantage appeared to diminish after that day. Overall, it is plausible that cassiosomes in the light eventually become limited in some other essential nutrients (e.g., nitrogen).

At the same time, the surprising survival of most cassiosomes in both treatments for two months suggests that cassiosomes do not entirely rely on symbiont photosynthesis for their carbon requirements. The abundance of lipid droplets visualized by SEM in the amoebocytes and the epidermis, coupled with a carbon-rich mesoglea core **(Figure 3A)**, may represent another main source of carbon that helped sustain viability. While changes in cassiosome size and appearance occurred with time, as previously described in *C. xamachana*, their survival in this study considerably exceeded the 10 days reported previously (17). In our study, the *in vitro* conditions (a stable and microbially-depleted environment) and high initial energy reserves (reflected in the abundance of lipid droplets) inherited from their regularly fed medusae of origin may have contributed to the long-term survival of the cassiosomes. However, our results highlight the surprisingly long-lived metabolic capacity of these small autonomous tissue structures. Further studies may investigate the survival and stinging capacity of cassiosomes over time in more natural environments.

### Ecological relevance

Our study indicates that the presence of symbiotic dinoflagellates in cassiosomes can increase their autonomous lifetime in the water column.

In this context, Anthony et al. (18) previously reported that the presence of dinoflagellates in cassiosomes of *C. ornata* differed between locations, and suggested that this difference could reflect different levels of investments in heterotrophic feeding. The differences in dinoflagellates abundance in cassiosomes could be due to differences in host dinoflagellate population densities and/or in the frequency of cassiosome release relative to the algal symbiont division rate. In any case, our results suggest that the presence of dinoflagellates in cassiosome may enhance their autonomous life span, thereby indirectly enhancing the heterotrophic feeding capacities of *Cassiopea* medusae.

Considering the importance of the algal symbionts in the life cycle of *Cassiopea*, and the specificity of this symbiotic interaction (38–41), it is surprising that *Cassiopea* produces only aposymbiotic larvae, which requires horizontal acquisition of dinoflagellates and selection of homologous symbionts by the polyp (42). In this context, it is possible that cassiosomes might constitute an environmental ‘reservoir’ of homologous symbionts in the environment, thereby facilitating symbiotic establishment for the newly formed polyps.

### Cassiosomes, a miniaturized model system for the cnidarian-Symbiodiniaceae symbiosis?

This study showed that early-released cassiosomes are metabolically active miniaturized holobionts that can effectively assimilate, exchange, and recycle nutrients autonomously. In particular, the symbiotic interface of the amoebocyte-dinoflagellate association seems to behave in a manner similar to other photosymbiotic cnidarians (such as corals), making cassiosomes a powerful laboratory model system for cell-to-cell symbiotic interactions (cell recognition, nutritional exchanges, etc). Their year-round availability and high abundance, easy collection process, and their simplified structural organization may in many situations prove advantageous over symbiotic polyps, larvae, or entire specimens of cnidaria. Their small size falls well within the technical limits of pristine vitrification by high-pressure freezing (around 200 μm, (43)) making them particularly suitable for in-depth characterization of cellular ultrastructure (e.g. using cryo-SEM). Studies of symbiotic interactions and of the protein composition of the symbiosome (i.e., with whole-mount immunolabeling experiments) also seem possible within these interesting ‘tissue balls’. Thus, even though the observed range in cassiosome shape and size might be a source of experimental variability, the preservation of symbiotic nutritional exchanges in these small cellular structures makes early released cassiosomes an attractive and powerful new miniaturized model system for the detailed study of the interface and machinery of cnidarian photosymbiosis.

## Conflicts of interest

None declared.

## Data availability

All raw data associated with this study have been deposited in the zenodo.org repository: https://doi.org/10.5281/zenodo.8038734.

## Author contribution statement

GT, NHL, GBP, CP, AM, NR conceived the experiment. GT, NHL, CP, NR performed the experiments. GT, GBP, CP, CMO, NR acquired and analyzed the data. GT wrote the first draft of the manuscript. All authors contributed to reviewing and revising the manuscript.

## Acknowledgements

The authors would like to thank C. Genoud, J. Daraspe and D. De Bellis for their advice on sample preparation and EM observations. A. Daley and the ANOM lab team at the University of Lausanne are thanked for sharing their aquarium facility. GT, GBP, AM and NR were supported by the Swiss National Science Foundation grants 200021_179092 and 212614. CP was supported by the Junior Professorship Grant ‘A connected underwater world’ awarded by the French National Research Agency and an associated start-up grant by the French National Centre for Scientific Research (CNRS).

## Notes

https://doi.org/10.5281/zenodo.8038734

### Competing Interest Statement

The authors have declared no competing interest.

